# Insufficient evidence for natural selection associated with the Black Death

**DOI:** 10.1101/2023.03.14.532615

**Authors:** Alison R. Barton, Cindy G. Santander, Pontus Skoglund, Ida Moltke, David Reich, Iain Mathieson

## Abstract

Klunk et al. analyzed ancient DNA data from individuals in London and Denmark before, during and after the Black Death [1], and argued that allele frequency changes at immune genes were too large to be produced by random genetic drift and thus must reflect natural selection. They also identified four specific variants that they claimed show evidence of selection including at *ERAP2*, for which they estimate a selection coefficient of 0.39–several times larger than any selection coefficient on a common human variant reported to date. Here we show that these claims are unsupported for four reasons. First, the signal of enrichment of large allele frequency changes in immune genes comparing people in London before and after the Black Death disappears after an appropriate randomization test is carried out: the *P* value increases by ten orders of magnitude and is no longer significant. Second, a technical error in the estimation of allele frequencies means that none of the four originally reported loci actually pass the filtering thresholds. Third, the filtering thresholds do not adequately correct for multiple testing. Finally, in the case of the *ERAP2* variant rs2549794, which Klunk et al. show experimentally may be associated with a host interaction with *Y. pestis*, we find no evidence of significant frequency change either in the data that Klunk et al. report, or in published data spanning 2,000 years. While it remains plausible that immune genes were subject to natural selection during the Black Death, the magnitude of this selection and which specific genes may have been affected remains unknown.

## 1. The signal of enrichment of unusual frequency changes at immune genes, a primary motivation for the search for individual signals, is a statistical artifact not driven by evolution

A primary motivation for Klunk et al.’s search for individual variants that changed in frequency after the Black Death is the finding that in a survey of thousands of variants in immune genes, there was highly significant enrichment of large *F_ST_* values compared to a panel of putatively neutral genomic regions (*P* < 10^-11^). However, when we randomly permuted which samples were labeled as pre- and post-Black Death, such that there should be no enrichment signal due to differences between time periods (**Methods**), we continued to find an extremely strong signal with inflated *P* values for the binomial test for enrichment over the 99th percentile of variants with minor allele frequency greater than 10% (**Figure 1A**). The *P* values do not follow a null expectation, and instead the χ^2^ values had an average inflation factor of 11.80 (**Figure 1B**). In particular, 7% of the permutations produce a *P* value more significant that that observed in the original paper (*P* = 7.9×10^-12^), corresponding to an increase of ten orders of magnitude to a non-significant level (*P* > 0.05). This is the pattern expected from a data artifact related to systematic coverage differences between the sets of immune and neutral loci (mean coverage at GWAS loci 7.1x, at exonic loci 2.9x and at neutral loci 8.3x), and not what would be expected if the signal was due to natural selection. We also carried out an independent set of 100 simulations, taking genome-wide sequencing data from present-day British and North American individuals with Northern European ancestry [2], and randomly assigning individuals to being before and after the Black Death while down-sampling the sequence data to approximately match the coverage of the individuals used in the actual published study on a per-site basis. This analysis showed enrichment of large *F_ST_* values at immune loci with an average inflation factor of 68.08 of the χ^2^ values (**Methods, Figure 1C,D**) and 39% of simulation replicates resulted in *P* values more significant than observed in the original paper. Taken together, these results demonstrate that differences in coverage at the tested sites can produce *P* values that do not follow a null expectation and can thus generate a spurious signal of enrichment similar to that reported by Klunk et al.

**Figure 1:**
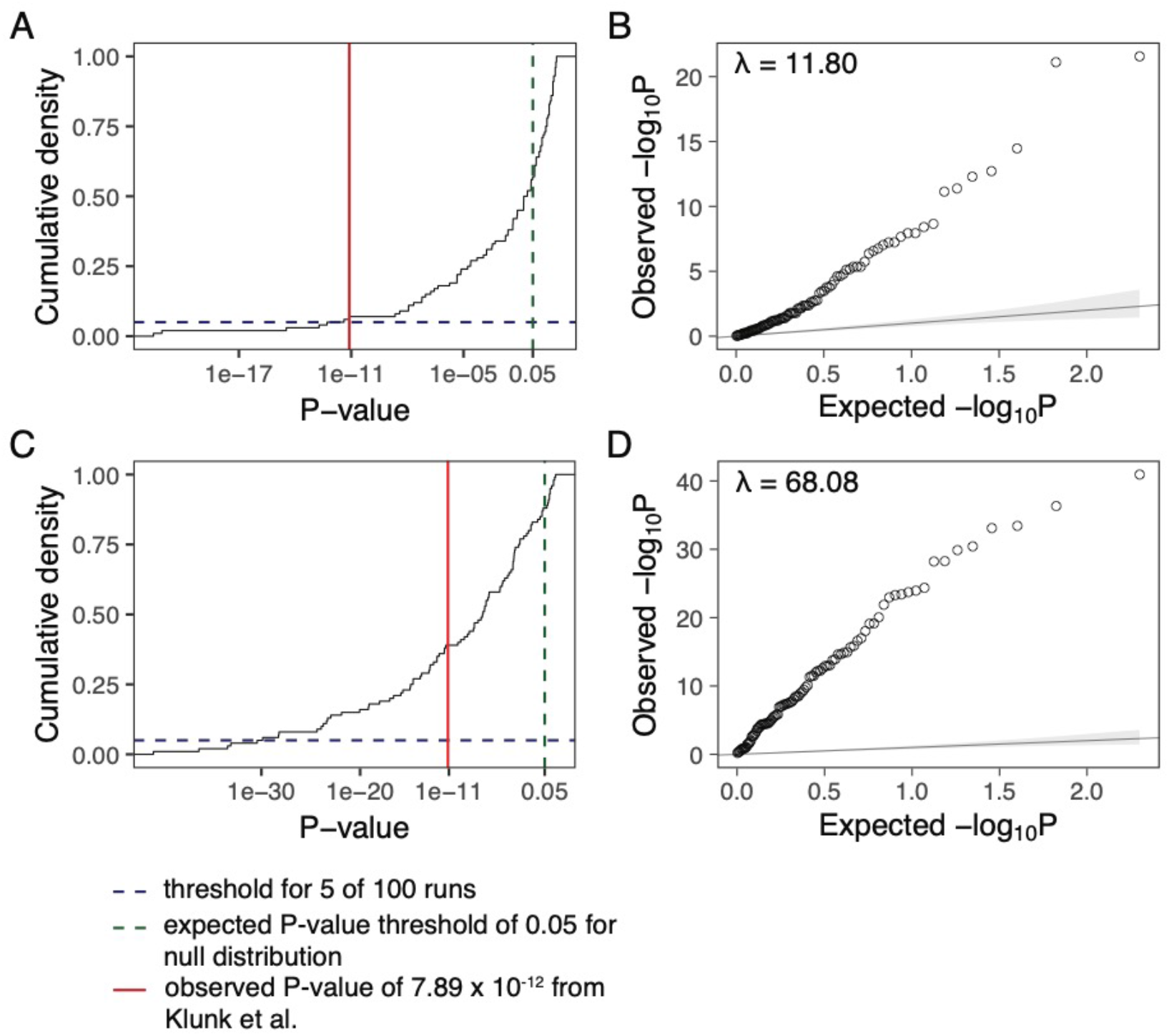
Reported enrichment persists even after permutation to remove any signal that could be due to the Black Death. **A:** *For 100 permutations of the pre- and post-Black Death labels, the observed p-values are plotted in a cumulative density plot for the 99^th^ percentile of enrichment of variants with MAF > 10%. The blue dashed line indicates a cumulative count of 5 of 100 runs, the green dashed line indicates the expected significance threshold of 0.05, and the red solid line shows the p-value obtained in the original paper*. **B:** *A q-q plot for the same p-values as in* **A** *showing the inflation over the expected null distribution of p-values*. **C:** *For 100 iterations approximately matching the sample size and coverage from the original study by downsampling a subset of samples from the 1000 Genomes Project, the observed p-values are plotted in a cumulative density plot for the 99^th^ percentile of enrichment of variants with MAF > 10%. Lines represent the same values as in* **A**. **D:** *A q-q plot for the same p-values as in* **D** *showing the inflation over the expected null distribution of p-values*.

## 2. Allele frequencies are estimated incorrectly and consequently biased; computed correctly, none of the four reported loci pass the filtering thresholds

Because it is difficult to make genotype calls from low coverage data, it is common to estimate allele frequencies based on genotype likelihoods. Typically, this is done using a maximum likelihood approach [3]. With ancient DNA, it is common to use pseudohaploid calls for robustness [4], although maximum likelihood approaches based on genotype likelihoods can also be used [5]. However, Klunk et al. do not do this and instead estimate allele frequencies using the genotype likelihoods as follows: “we calculated the expected number of alternate alleles as the likelihood the individual is heterozygous plus 2x the likelihood the individual is homozygous alternate.” This procedure is incorrect. Genotype likelihoods are the probability of the data conditional on the genotype *P*(*data|genotype*) but the approach of Klunk et al. treats them as though they were the probability of the genotype conditional on the data *P*(*genotype|data*). Their expectation is mathematically equivalent to a posterior mean with a prior that all three genotypes are equally likely. This will produce allele frequency estimates that are biased towards the prior mean of 0.5 and the extent of the bias will depend on coverage (because the posterior information depends on the read depth; **Figure 2A-C**). Thus, estimated differences in allele frequency across time periods can be driven by differences in coverage, rather than real changes in allele frequency (mean coverage in London pre-Black Death 6.8x, during Black Death 7.5x, post-Black Death 6.0x). Using the genotype likelihoods reported by Klunk et al., we recalculated allele frequencies using maximum likelihood and applied their filtering criteria (**Methods**). None of the four originally reported loci pass the thresholds used by Klunk et al. with the correctly estimated allele frequencies (one other locus does, but does not pass a Bonferroni-corrected threshold; **Figure 2D**). Rerunning the enrichment analysis of the coverage-matched data described in the previous section with this maximum likelihood allele frequency estimates still resulted in inflated *P* values indicating that this issue is separate to the one described above. Finally, we note that in order to avoid false positive calls due to damage most ancient DNA studies have either restricted to known polymorphic sites, restricted to transversions, or used Uracil-DNA glycosylase. Klunk et al. do not use any of these strategies. Of the 22,868 variants with MAF>5% that they analyzed, the transition/transversion ratio is around 19 (compared to an expectation of 2-3 for genomic data) and only 4,458 appear in the ~96 million UK Biobank SNP imputation set [6] which would be expected to include almost all sites with MAF>5% in the ancient populations. This suggests that the majority of the variants analyzed by Klunk et al. are false positives, likely due to ancient DNA damage.

**Figure 2:**
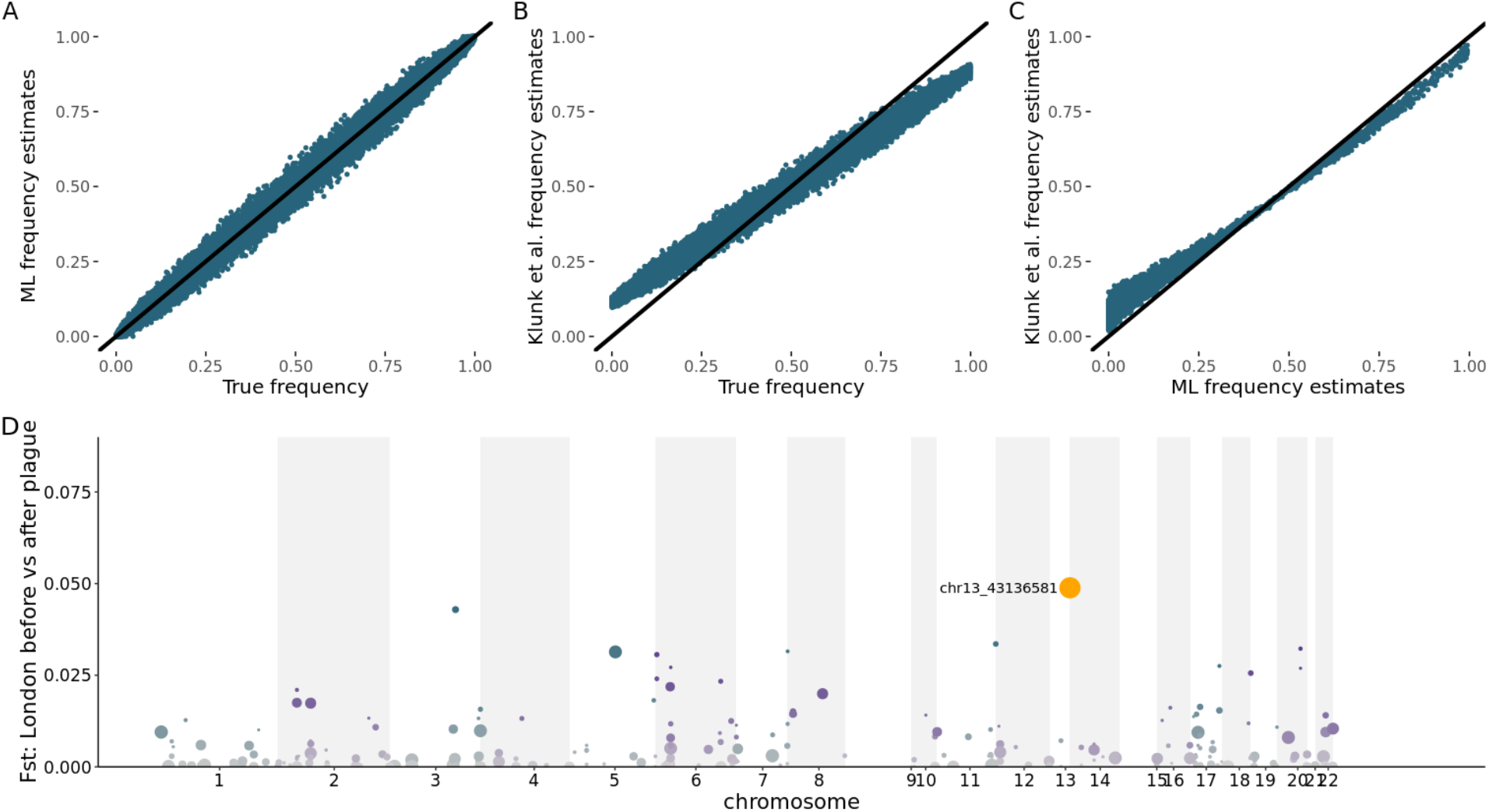
Bias in allele frequency estimates based on genotype likelihoods. **A**: *True values against unbiased maximum likelihood (ML) estimates for simulated read data with an average of 5x coverage simulated for 200 individuals at 30,000 SNPs to match the data observed in Klunk et al*. **B:** *True values against biased estimates computed with the Klunk et al. approach for simulated read data with an average of 5x coverage simulated for 200 individuals at 30,000 SNPs*. **C:** *Maximum likelihood (ML) estimates against Klunk et al. estimates for 31,799 SNPs included in the analysis with no minimum allele frequency threshold*. **D:** *Manhattan plot for F_ST_ scan of loci that pass the Klink et al. filtering criteria using maximum likelihood estimates of allele frequencies (equivalent to Figure 2C of Klunk et al.*).

## 3. Filtering thresholds do not appropriately correct for multiple testing

In order to control the experiment-wide false-positive rate, genome-wide scans must correct for the large number of statistical tests performed [7]. Klunk et al. do not apply such corrections, instead using an *ad hoc* filtering strategy with no clear statistical justification. Specifically, their filtering strategy requires variants to be in the top 5% of the *F_ST_* distribution in London and the top 10% in Denmark. Additionally, the variants are required to first increase then decrease (or vice-versa) in frequency in London and to move in the same direction in London and Denmark, when comparing pre-to post-Black Death samples. We simulated variants under a null model of identical allele frequencies in all populations using the same sample sizes as Klunk et al. (optimistically assuming diploid coverage at all loci) and find that a proportion 1.4×10^-4^ of variants pass these filters. For comparison, a conventional Bonferroni-corrected significance threshold would be 0.05/3293 = 1.5×10^-5^, approximately ten times smaller. Klunk et al. tested 3293 variants and should therefore expect 3293×1.4×10^-4^ = 0.46 variants to pass, if there were no difference in allele frequencies across populations. This is consistent with the one that we find when correctly estimating allele frequencies. Furthermore, many ancient DNA studies have required multiple linked variants to exceed the significance threshold in order to remove spurious outliers driven by genotyping errors [5, 8]-a requirement that is also standard in genome-wide association studies. Due to the sparse genotyping, Klunk et al. cannot apply this additional filter.

## 4. No evidence of natural selection at *ERAP2* over the past 2000 years

Klunk et al. show that almost all of their candidate immune genes are differentially expressed in response to stimulation by multiple pathogens including *Y. pestis*. They additionally show that their reported top SNP at *ERAP2* (rs2549794) is associated with a differential response to *Y. pestis* stimulation. However, this does not prove that the rs2549794 C allele is protective against infection, nor does it provide evidence of natural selection at the SNP – which can only be proven by a statistical analysis showing a change in population frequency due to Black Death exposure. However, the sample sizes in the study are so small that the uncertainty in *F_ST_* estimates is too large to infer selection. If there were in reality no difference in allele frequency, the probability of observing an *F_ST_* at least as high as the value of 0.0247 observed for rs2549794 with the available pre-vs post-Black Death London sample sizes and perfect genotype information would be approximately *P* = 0.067 (**Methods, Figure 3A**). This is likely an underestimate since in reality low coverage data, genetic drift and reference bias [9] could lead to inflated *F_ST_* values. The large point estimate of the selection coefficient—approximately five times larger than that at the lactase persistence allele [10] which is the largest selection coefficient at a common variant documented in humans to date—is likely simply a consequence of the fact that candidate variants were required to have large estimated *F_ST_* values and is not an independent piece of evidence for selection (**Figure 3B**).

**Figure 3:**
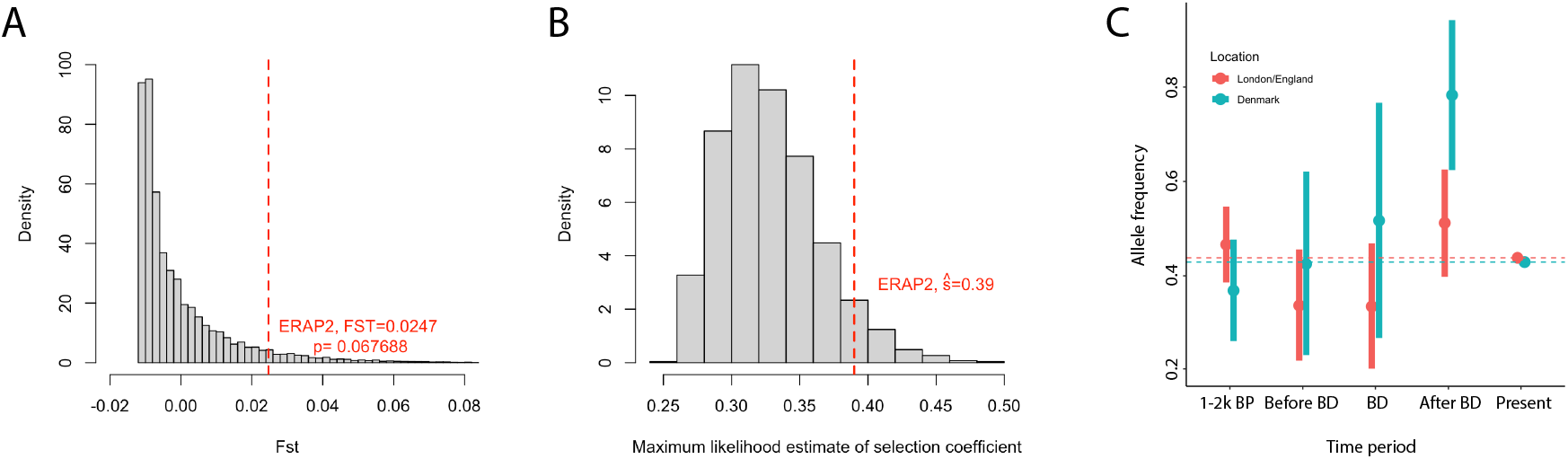
No evidence of selection at rs2549794. **A:** *Histogram of simulated F_ST_ values for two samples of sizes 38 and 63 diploid individuals from two populations with identical allele frequency of 0.438. Dashed red line shows reported value for rs2549794*. **B:** *Distribution of estimated selection coefficients, conditional on passing F_ST_ and directionality filters, under a null model of identical allele frequency of 0.438 in all populations assuming diploid coverage in all individuals*. **C:** *Estimated frequencies of rs2549794 in the three time points from Klunk et al. plus the periods 1-2K BP and the present day. Dashed lines show present-day frequencies and error bars show approximate 95% confidence intervals which overlap the present-day frequency for all time points except post-Black Death Denmark*.

We also examined the allele frequencies of rs2549794 in ancient and present-day populations from the United Kingdom [11–16] and Denmark [16, 17] to study whether there was any evidence for shifts in allele frequencies (**Methods**). The C allele frequency in present-day people with “British/Irish” ancestry from UK Biobank is 0.438 (N=264,261 alleles), not significantly different compared to ancient individuals from England dated between 2000 and 1000 BP (frequency=0.466, N=146 alleles, χ^2^ *P* = 0.55). Similarly, the frequency in present-day Denmark (0.429, N= 100,528 alleles) is not significantly different than Denmark 2000-1000 BP (0.368, N=76 alleles, *P* = 0.34). There is therefore no evidence that the frequency of rs2549794 has changed in either England or Denmark over the past 2,000 years (**Figure 3C**). Finally, we note that this SNP shows no evidence of selection in other scans that attempt to detect adaptation in Britain in this time period using a range of different data and methods. These include a scan based on time series of ancient DNA (rs2549794 empirical *P* = 0.18) [10], the singleton density score based on high coverage sequence data (rs2549794 Z=1.33, *P* = 0.09) [18] and a scan based on estimated coalescent times in UK Biobank (minimum *P* value within 1Mb 0.40) [19]. The locus also failed to replicate in two independent studies of ancient genomes from before and after the Black Death in Britain and Norway [9, 20].

## Conclusion

Inadequate correction for multiple testing, implausibly large estimated effects due to small sample sizes, and *post hoc* rationalization for marginal associations are precisely the issues that led to the development of clear statistical standards for genome-wide scans. Even ignoring technical issues with the allele frequency estimation and systematic differences in coverage between loci, the evidence presented by Klunk et al. does not meet these standards.

## Methods

### Permutation test for enrichment at immune loci

To test whether the observed enrichment of large *F_ST_* values at immune loci was due to real signals of differentiation or to technical artifacts related to the samples and targeted loci, we performed a permutation analysis. We randomly shuffled the labels for the London pre-Black Death and London post-Black Death samples such that for every run of the permutation test, the new cohort for each time point consisted of random individuals drawn from the union of the original time points without replacement. We also preserved the number of samples in each time point included for each target panel (GWAS loci, exon loci, and neutral loci). We performed this sampling 100 times, and ran each set of samples through the original Klunk et al. allele frequency estimation and enrichment pipeline, testing for enrichment at a 99% threshold for variants with MAF > 0.10.

### Testing whether differences in coverage can lead to apparent enrichment at immune loci

The differences in the mean coverage of the three target panels suggested a possible cause of the observed enrichment of high *F_ST_* values at the immune loci. To test this, we replicated 100 datasets with approximately matched coverage to the samples sequenced in Klunk et al. and ran them through their original pipeline. For each run, we randomly sampled 206 of the 270 GBR and CEPH samples from the 1000 Genomes Project [2] sequenced to ~30x to match each of the samples in the Klunk et al. study that were not dropped from the final analysis due to high missingness. We computed read depths using “samtools depth” [21] at all sites tested by Klunk et al. and present in the UK Biobank v3 imputation panel [6] for both datasets using the UCSC browser version of LiftOver to convert from hg19 sites to GRCh38 [22]. Next, we used the “samtools view -s” option to down-sample each 1000 Genomes alignment to the fraction of reads present in the original sample. Where the Klunk et al. read depth exceeded that in the 1000 Genomes sample read depth, we retained all reads. We then ran these down-sampled alignments through HaplotypeCaller to produce gvcfs with calls at each of the sites and then subsequently joint called again with GATK HaplotypeCaller [23] across all 207 simulated samples, in both instances using the same parameters as those in the original pipeline, again using the original allele frequency estimator. Finally, we used these calls as input for the original Klunk et al. pipeline. The only alteration to this method was that, because of the reduced number of sites compared to the original paper, the same samples were dropped or retained as in the original study regardless of their missingness in the new dataset. In addition to the results reported above, the enrichment of 99% for variants with MAF > 0.30 reported in Klunk et al. showed inflation of chi-squared statistics of 67.92, and 23% of the *P* values were more significant than that reported in the original study (*P*=1.16×10^-14^). When using a maximum likelihood method to calculate allele frequencies rather than the original method, the enrichment of 99% for variants with MAF > 0.10 resulted in 50% of *P* values more significant than that calculated for the original data with an updated maximum likelihood estimator (*P*=2.09×10^-16^) and an inflation of χ^2^ statistics of 147.66. Additionally, for the enrichment of 99% for variants with MAF > 0.30, 31% of *P* values were more significant than that calculated for the original data with an updated maximum likelihood estimator (*P*=1.64×10^-13^), and there was an inflation of χ^2^ statistics of 68.81.

### Maximum likelihood estimation of allele frequencies

Let the genotype likelihoods for individual *i=1…n* be given by 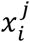 for genotypes *j=0,1,2*. Then the maximum likelihood estimate of the allele frequency *p* is:

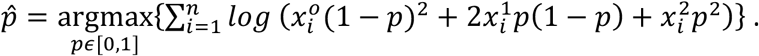

We implemented the maximum likelihood estimator by numerically maximizing the expression above using the genotype likelihoods reported by Klunk et al. In contrast, the Klunk et al. estimator is 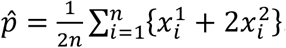.

### Evaluation of filtering threshold under a null model

We simulated 1 million observations of genotype data from populations with identical allele frequencies ranging from 0.1 to 0.5 in intervals of 0.1 with sample sizes corresponding to the cohort sizes of the London and Denmark samples. We then calculated *F_ST_* using the Weir and Cockerham estimator [24] and counted the proportion of observations that passed the reported filtering threshold, averaged across all allele frequencies.

### Selection coefficients for variants that pass the filtering threshold under a null model

*We* simulated genotype data for 10 million variants with observations from both London and Denmark using the same sample sizes as in Klunk et al. and assuming a frequency of 0.438 across both populations and all time points. We then applied the *F_ST_* and directionality filters from Klunk et al. to these variants and for the 1328 variants that passed the filters we applied the selection coefficient estimation code made available by Klunk et al.

### Allele frequencies and differentiation at rs2549794

We looked up ancient and present-day allele frequencies from the following sources:

1. Ancient England, from the Allen Ancient DNA Resource v54.1 (AADR) (https://reich.hms.harvard.edu/allen-ancient-dna-resource-aadr-downloadable-genotypes-present-day-and-ancient-dna-data), we extracted all individuals with “Locality” entry starting with “England” dated to between 2000 and 1000 BP [11, 13–16].
2. Ancient Denmark, from the AADR, we extracted all individuals with “Political Entity” equal to “Denmark” dated to between 2000 and 1000 BP [16].
3. Present-day England, we looked up the allele frequencies in the “British/Irish ancestry” subset of whole-genome sequenced UK Biobank participants in the deCAF browser (https://decaf.decode.com) [12].
4. Present-day Denmark, we looked up allele frequencies in the iPsych study [17].

For these observations, we estimated standard deviation of the allele frequencies using the Gaussian approximation to the binomial distribution. For the observations from Klunk et al. we estimated standard deviations of the maximum likelihood allele frequency estimates by bootstrapping with 10,000 replicates. We plot approximate 95% confidence intervals defined as ±1.96 estimated standard deviations.

## Data availability

No new data were generated for this publication. The sequence data from Klunk et al. were obtained from the authors. High coverage 1000 genomes data was obtained from https://www.internationalgenome.org/data-portal/data-collection/30x-grch38. Genotype data of other ancient samples was obtained from https://reich.hms.harvard.edu/allen-ancient-dna-resource-aadr-downloadable-genotypes-present-day-and-ancient-dna-data.

## Code availability

Code to replicate the analyses described here is available at https://github.com/arbarton/Klunk_matters_arising/

## Acknowledgments

PS was supported by the European Molecular Biology Organisation, the Vallee Foundation, the European Research Council (grant no. 852558), the Wellcome Trust (217223/Z/19/Z), and Francis Crick Institute core funding (FC001595) from Cancer Research UK, the UK Medical Research Council, and the Wellcome Trust. CGS and IMo were funded by a European Research Council starting grant awarded to IMo (ERC-2018-STG-804679). IMo was also supported by a Villum Fonden Young Investigator grant (project no. 19114). DR was supported by National Institutes of Health grant HG012287; John Templeton Foundation (grant 61220); the Allen Discovery Center program, a Paul G. Allen Frontiers Group advised program of the Paul G. Allen Family Foundation; and the Howard Hughes Medical Institute. IMa was supported by the National Institute of General Medical Sciences (GM133708). The content is solely the responsibility of the authors and does not necessarily represent the official views of the National Institutes of Health or other funding agencies.

## References

1. Klunk, J., et al., Evolution of immune genes is associated with the Black Death. Nature, 2022. 611(7935): p. 312–319.

2. Byrska-Bishop, M., et al., High-coverage whole-genome sequencing of the expanded 1000 Genomes Project cohort including 602 trios. Cell, 2022. 185(18): p. 3426–3440 e19.

3. Kim, S.Y., et al., Estimation of allele frequency and association mapping using next-generation sequencing data. BMC Bioinformatics, 2011. 12: p. 231.

4. Green, R.E., et al., A draft sequence of the Neandertal genome. Science, 2010. 328(5979): p. 710–722.

5. Mathieson, I., et al., Genome-wide patterns of selection in 230 ancient Eurasians. Nature, 2015. 528(7583): p. 499–503.

6. Bycroft, C., et al., The UK Biobank resource with deep phenotyping and genomic data. Nature, 2018. 562(7726): p. 203–209.

7. Risch, N. and K. Merikangas, The future of genetic studies of complex human diseases. Science, 1996. 273(5281): p. 1516–7.

8. Bergstrom, A., et al., Grey wolf genomic history reveals a dual ancestry of dogs. Nature, 2022. 607(7918): p. 313–320.

9. Gopalakrishnan, S., et al., The population genomic legacy of the second plague pandemic. Curr Biol, 2022. 32(21): p. 4743–4751 e6.

10. Mathieson, I. and J. Terhorst, Direct detection of natural selection in Bronze Age Britain. Genome Res, 2022. 32(11-12): p. 2057–2067.

11. Martiniano, R., et al., Genomic signals of migration and continuity in Britain before the Anglo-Saxons. Nature communications, 2016. 7: p. 10326.

12. Halldorsson, B.V., et al., The sequences of 150,119 genomes in the UK Biobank. Nature, 2022. 607(7920): p. 732–740.

13. Schiffels, S., et al., Iron age and Anglo-Saxon genomes from East England reveal British migration history. Nature communications, 2016. 7: p. 10408.

14. Patterson, N., et al., Large-scale migration into Britain during the Middle to Late Bronze Age. Nature, 2022. 601(7894): p. 588–594.

15. Gretzinger, J., et al., The Anglo-Saxon migration and the formation of the early English gene pool. Nature, 2022. 610(7930): p. 112–119.

16. Margaryan, A., et al., Population genomics of the Viking world. Nature, 2020. 585(7825): p. 390–396.

17. Erlangsen, A., et al., Genetics of suicide attempts in individuals with and without mental disorders: a population-based genome-wide association study. Mol Psychiatry, 2020. 25(10): p. 2410–2421.

18. Field, Y., et al., Detection of human adaptation during the past 2000 years. Science, 2016. 354(6313): p. 760–764.

19. Nait Saada, J., et al., Identity-by-descent detection across 487,409 British samples reveals fine scale population structure and ultra-rare variant associations. Nat Commun, 2020. 11(1): p. 6130.

20. Hui, R., et al., Medieval social landscape through the genetic history of Cambridgeshire before and after the Black Death. bioRxiv, 2022: p. doi: https://doi.org/10.1101/2023.03.03.531048.

21. Danecek, P., et al., Twelve years of SAMtools and BCFtools. Gigascience, 2021. 10(2).

22. Nassar, L.R., et al., The UCSC Genome Browser database: 2023 update. Nucleic Acids Research, 2022. 51(D1): p. D1188–D1195.

23. Poplin, R., et al., Scaling accurate genetic variant discovery to tens of thousands of samples. bioRxiv, 2017: p. DOI: 10.1101/201178.

24. Weir, B.S. and C.C. Cockerham, Estimating F-Statistics for the Analysis of Population Structure. Evolution, 1984. 38(6): p. 1358–1370.

